# FBXW7 alleviates c-MYC repression of pyruvate carboxylase to support metabolic flexibility

**DOI:** 10.1101/2024.10.24.619388

**Authors:** Miriam Lisci, Fanny Vericel, Hector Gallant-Ayala, Julijana Ivanisevic, Alexis A. Jourdain

## Abstract

Metabolic flexibility, or the ability to adapt to environmental fluctuations, is key to the survival and growth of all living organisms. In mammals, the pathways supporting cell proliferation in nutrient-limiting conditions have not been fully elucidated, although cancers are known to display metabolic dependencies that can be targeted for therapy. Here, we combine systematic nutrient and genome-wide CRISPR/Cas9 screening to provide a comprehensive map of the signaling and metabolic pathways that support cell proliferation in glutamine-limited conditions. We focus on pyruvate anaplerosis and discover a mechanism by which the tumor suppressor FBXW7 controls a MYC-dependent cluster of epigenetic repressors that bind the pyruvate carboxylase (*PC*) promoter, leading to histone deacetylation, reduced *PC* expression and glutamine addiction. Our work sheds light on the molecular mechanisms that support metabolic flexibility, and on the nutrients and pathways involved in glutamine dependency, a hallmark of several cancers.

**Graphical abstract:** 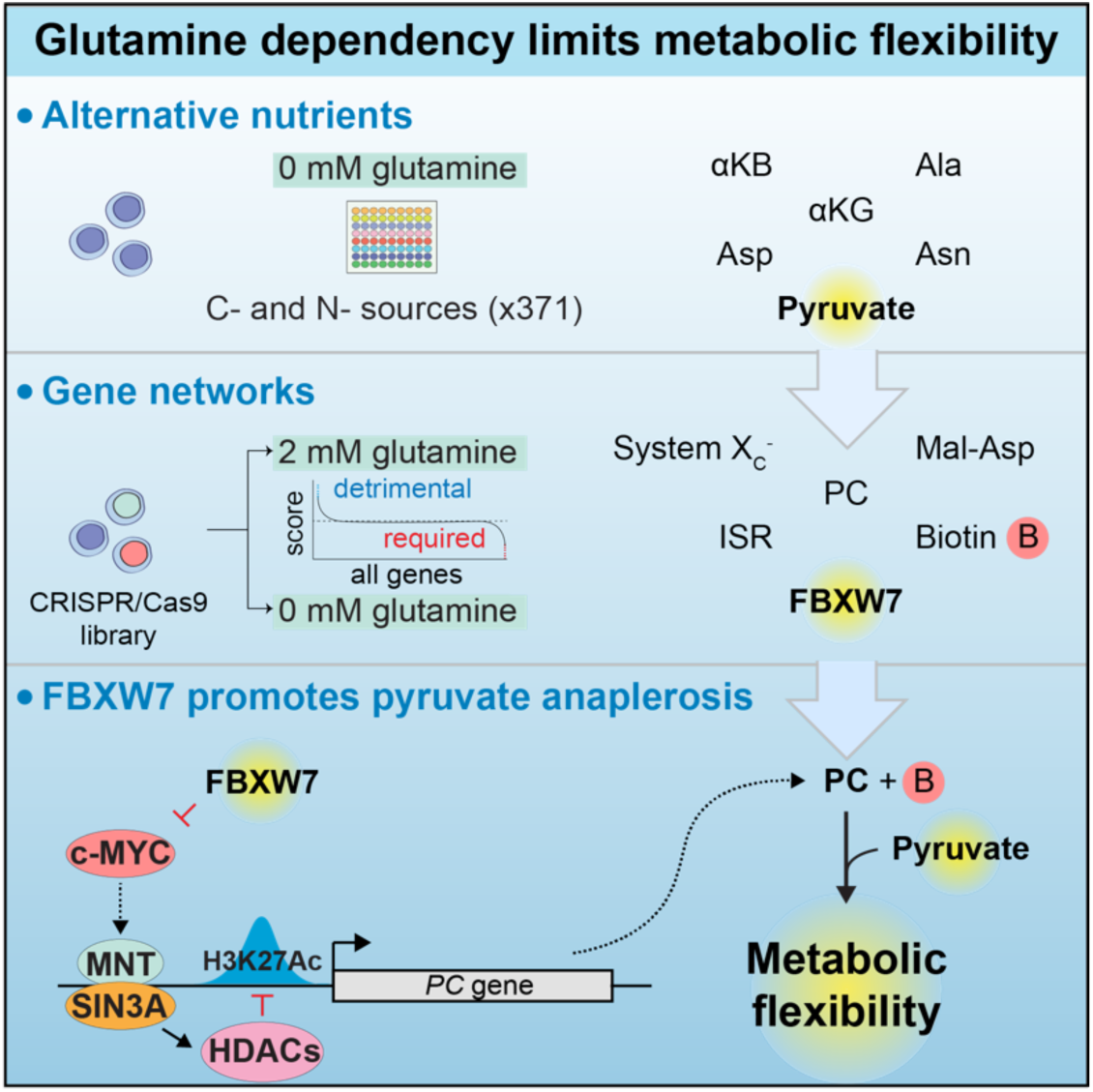

## Introduction

Mammalian cells have evolved to cope with fluctuating environments and conditions such as malnutrition, infection or tumorigenesis can further perturb access to nutrients, making metabolic flexibility crucial for ensuring viability^1^. Amino acids are among the nutrients that show the highest variation both in our diet and across tissues^2^, and among them glutamine is the most abundant amino acid in plasma and one of the most highly utilized amino acids in cells in culture^3,4^. Besides its role as an essential constituent of proteins, glutamine acts as a major contributor both in the tricarboxylic acid (TCA) cycle, where it participates in ATP generation and amino acids synthesis, and in nitrogen metabolism, serving as an amine donor in multiple metabolic reactions.

While glutamine is classically viewed as a non-essential amino acid, it has long been known that mammalian cells can become dependent on its extracellular availability^4^. This phenomenon, sometimes referred to as “glutamine addiction” has been observed both in cell culture and *in vivo*, and it is thought to be the result of high proliferation rates which exceed the capacity for its intracellular biosynthesis^5^. In tumors, glutamine addiction is mainly driven by the oncogene c-MYC (*MYC*), which promotes the expression of glutamine transporters and glutaminase^6–8^, thereby converting glutamine into glutamate and facilitating its entry into the TCA cycle. This process has been exploited for cancer therapy based on glutaminase inhibition, although with only limited success in pre-clinical trials^9^, partly due to the metabolic flexibility of cancer cells and the presence of alternative nutrients in the complex tumor microenvironment. A better understanding of the molecular mechanisms that promote cell survival in a glutamine-limited environment could help to identify novel approaches to limit cancer cell proliferation.

Earlier studies have highlighted certain nutrients and metabolic pathways that can sustain proliferation of glutamine-deprived cells, including pyruvate carboxylation, the malate-aspartate shuttle and asparagine synthesis^10–12^, as well as other pathways that become detrimental in the absence of glutamine, such as proline synthesis^13^. However, despite this important work, we still lack a global view of the nutrients and metabolic pathways involved in cell survival when glutamine is scarce. Here, we combine large-scale nutrient and genetic screens to provide a unified model of the metabolites and molecular pathways that sustain viability upon glutamine starvation. We further reveal an epigenetic mechanism whereby the tumor suppressor *FBXW7* alleviates glutamine dependency by promoting expression of pyruvate carboxylase (*PC*), thus favoring pyruvate anaplerosis and proliferation in nutrient-limiting conditions.

## Results

### Nutrient screening identifies molecules increasing proliferation in a glutamine-limited environment

We sought to explore the mechanisms that maintain proliferation during glutamine deprivation, and observed that glutamine depletion led to a rapid loss of proliferation and viability in K562 myelogenous leukemia cells, indicating strict dependency on glutamine availability (**Figure 1A, B; Figure S1A**). Reasoning that the cell culture medium represents only a fraction of the molecules available to cells *in vivo*, we designed a nutrient screen where we supplemented the medium with 371 carbon and nitrogen sources in order to identify nutrients able to compensate for glutamine depletion (**Figure 1C**). We found a wide range of metabolites able to sustain proliferation to various extents, including carboxylic acids such as α-ketoglutarate (α-KG) and α-ketobutyrate (α-KB), and amino acids chemically related to glutamine, some of which had previously been observed in the context of glutamine deprivation or glutaminase inhibition^12,14^ (**Figure 1D, E; Table S1**). In our screen, we found that pyruvate provided the most robust increase in proliferation in glutamine-deprived medium, a result that we could validate in additional cell lines (**Figure 1F; Figure S1B**).

**Figure 1.**
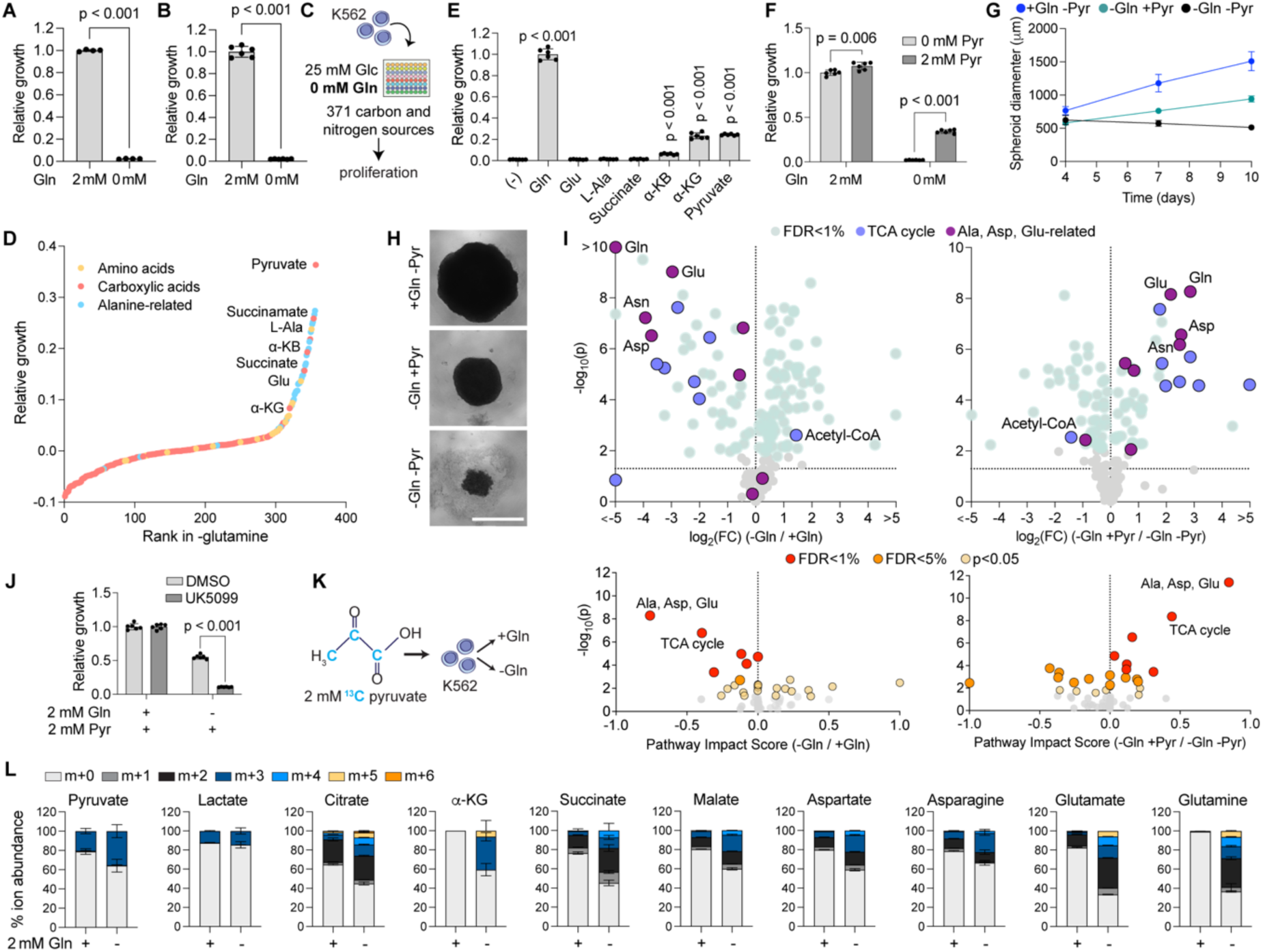
Nutrient screening ranks metabolites that allow proliferation of glutamine-deprived K562 cells. **(A, B)** Cell count-based (A) or Prestoblue (B) growth assays of K562 cells in medium with or without glutamine (Gln). p values: unpaired t-test. **(C)** Schematic representation of the nutrient-sensitized screen in high glucose (Glc) medium without glutamine. **(D)** S-plot of nutrients promoting proliferation in glutamine-deprived medium. Glutamine and glutamine-containing dipeptides not shown. **(E)** Validation of selected nutrients (all supplemented at 2 mM) in glutamine-starved cells. Glu: glutamate; Ala: alanine; α-KB: α-ketobutyrate; α-KG: α-ketoglutarate. p values: one-way ANOVA. **(F)** Growth assay in the presence of pyruvate (Pyr) with or without glutamine. p values: multiple unpaired t-tests. **(G)** Diameter of HEK293T spheroids cultured in medium supplemented with either glutamine or pyruvate. **(H)** Representative spheroid images on day 10 post-seeding. Scale bar = 1 mm. **(I)** Volcano plots showing metabolite abundance in glutamine-deprived vs glutamine-rich medium (top left) or by pyruvate supplementation in glutamine-free conditions (top right). TCA intermediates and metabolites related to alanine, glutamate, aspartate (Asp) and asparagine (Asn) are labeled. Bottom: MetaboAnalyst Pathway Analysis of significantly altered metabolites. **(J)** Proliferation assay in the absence or presence of the mitochondrial pyruvate carrier inhibitor (UK5099). p values: multiple unpaired t-tests. **(K)** ^13^C pyruvate tracing strategy. **(L)** Results of ^13^C pyruvate tracing in glutamine-rich (2 mM) and glutamine-deprived medium, showing relative fractions of isotopologues within each metabolite.

Since nutrient availability has been well-characterized as a limiting factor in solid tumors due to poor vascularization^15^ we tested our findings using a three-dimensional spheroid model to determine whether pyruvate could also increase cell proliferation in a more physiological model of glutamine deprivation. We grew HEK293T cells in medium supplemented with either glutamine or pyruvate and measured spheroid diameter at regular time intervals. As expected, we found that in the absence of glutamine spheroids failed to grow and developed morphological characteristics consistent with cell death, while growth of spheroids was restored in the presence of glutamine (**Figure 1G, H**). Furthermore, addition of pyruvate also significantly rescued both the growth and appearance of spheroids in glutamine-depleted medium, consistent with the results obtained using K562 cells.

We next examined which metabolic pathways were most affected by glutamine and pyruvate. In cells with certain mitochondrial defects, pyruvate and the enzyme prosthetic *Lb*NOX are able to correct the NADH/NAD^+^ ratio and restore cell growth^16,17^. However, we found that *Lb*NOX had only a minimal effect in our glutamine-restricted conditions, indicating that restoring the NADH/NAD^+^ ratio is not the primary effect of pyruvate under glutamine deprivation (**Figure S1C**). We therefore performed steady-state metabolomics following glutamine restriction and found that it severely decreased the abundance of all TCA cycle intermediates, as well as certain amino acids such as glutamate, aspartate and asparagine (**Figure 1I; Table S2)**. The levels of metabolites related to glycolysis, nucleotide synthesis, the pentose phosphate pathway, as well as other amino acids showed little to no variation in the absence of glutamine. We then repeated this experiment using medium supplemented with pyruvate and observed a significant rescue in all detected TCA cycle intermediates and in the same three amino acids (**Figure 1I; Table S3**), suggesting that the anaplerotic contribution of pyruvate to the mitochondrial TCA cycle increased during glutamine deprivation.

To test this hypothesis, we treated cells with the mitochondrial pyruvate carrier inhibitor UK5099 and observed that while cells grown in glutamine-rich medium did not show any decrease in proliferation, the growth advantage conferred by pyruvate supplementation in the absence of glutamine was almost completely abolished in the presence of this inhibitor (**Figure 1J**). Furthermore, to investigate a possible effect of pyruvate in anaplerosis we performed ^13^C pyruvate-based isotope labeling (**Figure 1K**) and found increased ^13^C incorporation in all detected TCA cycle intermediates in absence of glutamine, as well as in glutamate, aspartate, asparagine and in glutamine itself (**Figure 1L; Table S4**). Combining the results from our nutrient screen and stable isotope-assisted profiling experiment, we conclude that the contribution of pyruvate to the TCA cycle and to the synthesis of selected amino acids increases during glutamine deprivation to restore cell proliferation.

### Genetic screening reveals metabolic networks promoting pyruvate anaplerosis in absence of glutamine

We next designed a genome-wide screen aimed at identifying the molecular pathways able to maintain proliferation in glutamine-free, pyruvate-rich conditions. We infected K562 cells with the Brunello lentiviral genome-wide CRISPR/Cas9 library with 76,441 single-guide RNAs (sgRNAs) targeting 19,114 distinct human genes and 1,000 controls. After incubation for 10 days to allow for CRISPR/Cas9-mediated gene knockout, we transferred the cells to a DMEM-based medium containing glucose and pyruvate, either with or without glutamine. After 21 days we harvested the cells and performed next-generation sequencing to determine the abundance of sgRNAs in each condition. We analyzed the data using a Z score-based method^18^, highlighting genes that were either required (negative ΔZ score) or detrimental (positive ΔZ score) for growth of the glutamine-starved cells (**Figure 2A; Table S5)**.

**Figure 2.**
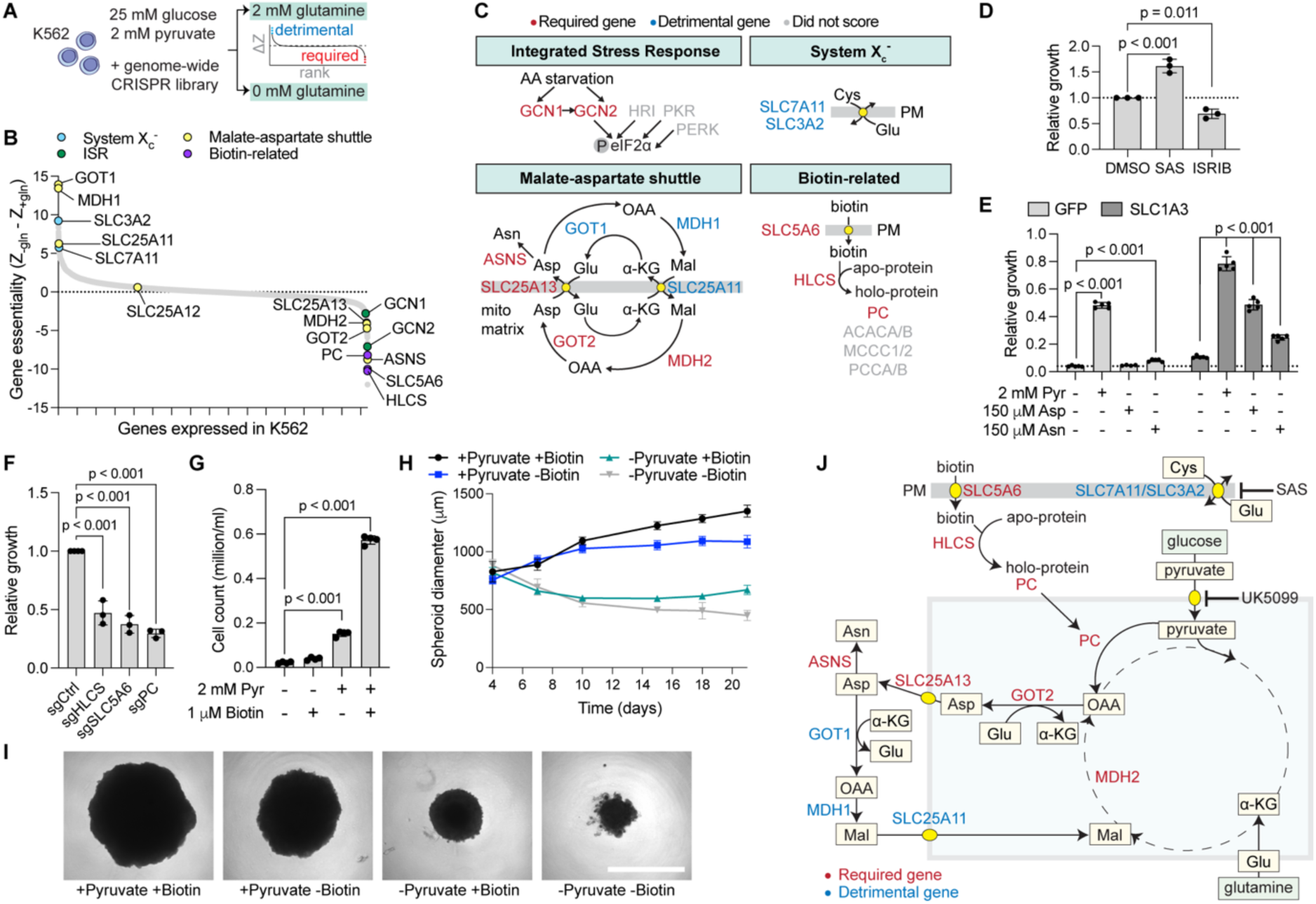
Genetic screening provides a unified model of pathways promoting pyruvate anaplerosis during glutamine starvation. **(A)** Genome-wide CRISPR/Cas9 screening strategy and analysis. **(B)** Essentiality score of genes promoting (ΔZ<0) or inhibiting (ΔZ>0) proliferation upon glutamine depletion. ISR: integrated stress response. **(C)** Overview of pathways required (ΔZ<2, red) or detrimental (ΔZ>2, blue) for growth in glutamine-limiting conditions. Genes below the selected thresholds are labeled as “did not score”. PM: plasma membrane. **(D)** Proliferation of glutamine-starved K562 cells treated with inhibitors of the X_c_^-^ antiporter (sulfasalazine, SAS) or the ISR (ISRIB). **(E)** Proliferation of glutamine-starved K562 cells expressing the plasma membrane amino acid transporter SLC1A3 or GFP as a control. Cells were grown in glutamine-deprived medium supplemented with pyruvate, aspartate or asparagine. **(F)** Proliferation of glutamine-starved cells depleted of biotin-related genes. All conditions were supplemented with 2 mM pyruvate. **(G)** Growth assay in glutamine-deprived medium upon addition of pyruvate, biotin or both. **(H)** Quantification of spheroid diameter of glutamine-restricted HEK293T supplemented with biotin, pyruvate or both. **(I)** Representative images for HEK293T spheroids grown in glutamine-deprived medium in the same conditions as in (H), 20 days post-seeding. Scale bar = 1 mm. **(J)** Unified model of metabolic networks related to proliferation in glutamine-deprived medium. All p values: one-way ANOVA.

Our screen highlighted several pathways known to participate in central carbon metabolism, including the malate-aspartate shuttle, the plasma membrane cystine/glutamate exchanger system X_c_^-^ biotin import and conjugation, pyruvate carboxylation, as well as signaling molecules from the integrated stress response (ISR) (**Figure 2B, C**). To validate the robustness of our approach, we confirmed experimentally that blocking ISR signaling downstream of the amino acid-sensing arm (genes *GCN1*, *GCN2*) using ISRIB, a drug that reverses the effect of eIF2α phosphorylation, was detrimental to cell growth in the absence of glutamine, while inhibition of the system X_c_^-^ (*SLC3A2, SLC7A11*) involved in glutamate extracellular export, was beneficial (**Figure 2C, D**). We also focused on a subset of genes involved in the malate-aspartate shuttle and found that our screen not only confirmed the requirement for the mitochondrial arm of the shuttle (*GOT2*, *MDH2, SLC25A13*)^14^, but it also revealed that deleting the cytosolic arm (*GOT1*, *MDH1, SLC25A11*), involved in bringing intermediates back to mitochondria, was beneficial in conditions of glutamine deprivation. This screen further confirmed an important role for the *ASNS* gene^11,12^, required for asparagine synthesis. These observations, along with earlier literature, strongly suggested an important role for aspartate synthesis and cytosolic export, as well as for synthesis of asparagine. We confirmed this hypothesis by providing these two amino acids to cells grown in glutamine-deprived medium and expressing *SLC1A3,* a plasma membrane transporter required for aspartate import^16^, upon which we observed a partial rescue in cell proliferation (**Figure 2E**).

Notably, our screen identified a requirement for biotin-related genes for the growth of glutamine-deprived cells. Biotin, also known as vitamin B7 or vitamin H, is the substrate for biotinylation, a post-translational modification that occurs in 4 different classes of mitochondrial carboxylases, one of which, pyruvate carboxylase (*PC*), was identified in our screen (**Figure 2B, C**). PC converts pyruvate and CO_2_ into oxaloacetate, and a beneficial role for pyruvate carboxylation in glutamine-limited environment has previously been noted^10^. Importantly, our ^13^C pyruvate tracer analysis showed increased fraction of the m+3 isotopologues of malate, aspartate and asparagine (**Figure 1L**), all of which are compatible with increased PC activity in glutamine-limiting conditions. Consistent with this, we confirmed experimentally that, in addition to *PC*, the biotin plasma membrane transporter (*SLC5A6*) and the biotin conjugating enzyme (*HLCS*) were both required for the proliferation of cells in the absence of glutamine (**Figure 2F**). We further validated these observations using biotin supplementation in glutamine-deprived cells and confirmed a requirement for both pyruvate and biotin to sustain K562 cell proliferation (**Figure 2G**) and HEK293T spheroid formation (**Figure 2H, I**).

Together, our genetic and nutrient screens provide a wide range of factors whose depletion is either deleterious, or in some cases beneficial, for the proliferation of cells in the absence of glutamine. Importantly, the genes and nutrients we have identified highlight interconnecting pathways and, together with earlier literature, point to a unified model where glutamine-restricted cells rely on import of pyruvate into mitochondria (**Figure 1J**) and its conversion to oxaloacetate by the biotin-dependent carboxylase PC, thereby supporting mitochondrial aspartate synthesis, its export via the malate-aspartate shuttle, and asparagine synthesis in the cytosol (**Figure 2J**).

### *FBXW7* supports PC expression and pyruvate anaplerosis

While most hits from our genetic screen are involved in central carbon metabolism, the gene that scored the highest in the absence of glutamine was the tumor suppressor *FBXW7* (**Figure 3A**). FBXW7 is a well-characterized substrate recognition component for the cytosolic SKP1-CUL1-F-box E3 ubiquitin ligase complex, involved in degradation of the protein products of oncogenes such as c-MYC, c-JUN, and cyclin E^19–22^.

**Figure 3.**
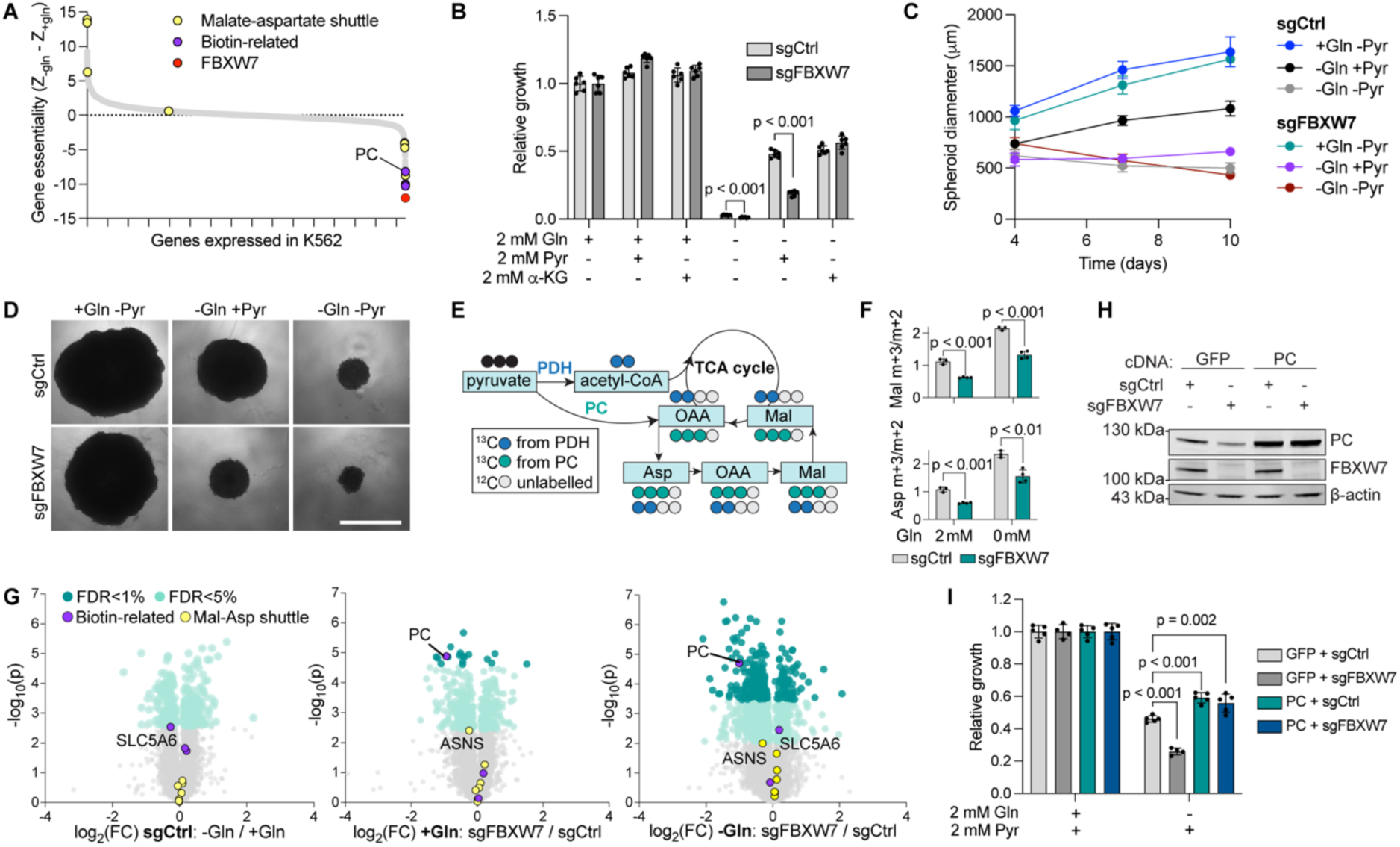
*FBXW7* mediates pyruvate anaplerosis by promoting PC expression. **(A)** Genome-wide CRISPR/Cas9 screen reveals *FBXW7* as the top hit required for proliferation upon glutamine depletion. **(B)** Growth assay of control (sgCtrl) and *FBXW7*-depleted cells (sgFBXW7) in media supplemented with glutamine, pyruvate or α-KG. p values: multiple unpaired t-tests. **(C)** Quantification of spheroid diameter for control and *FBXW7-*depleted HEK293T grown in media supplemented with either glutamine or pyruvate. **(D)** Representative images of spheroids cultured as in (C) on day 10 post-seeding. Scale bar = 1 mm. **(E)** Schematic representation of ^13^C pyruvate tracing for a subset of TCA intermediates. OAA: oxaloacetate; Mal: malate. **(F)** m+3/m+2 ratio of ^13^C pyruvate-derived carbons for malate and aspartate in control and *FBXW7*-depleted cells grown in glutamine-rich or -deprived medium. **(G)** Volcano plots showing differential protein expression in glutamine-rich (2 mM) versus glutamine-depleted medium (sgCtrl, left), and in *FBXW7*-depleted cells in glutamine-rich (center) and glutamine-deprived (right) medium, compared to sgCtrl. **(H)** Immunoblot showing PC and FBXW7 expression in control and *FBXW7*-depleted cells overexpressing *PC* cDNA, or a GFP control. **(I)** Growth assay of samples shown in (H) cultured in medium supplemented with pyruvate, and with or without glutamine. p values: one-way ANOVA.

Since a role for *FBXW7* in central carbon metabolism was unexpected, we depleted *FBXW7* in K562 cells using CRISPR/Cas9. We observed that while *FBXW7*-depleted cells showed no proliferation defects in glutamine-rich medium, they grew more slowly in the absence of glutamine, regardless of the presence of pyruvate in the medium (**Figure 3B**). To investigate whether this difference was specifically due to pyruvate metabolism, we supplemented the cell medium with α-KG and we found that this metabolite was able to rescue cell proliferation independently of the presence of *FBXW7*, pointing to a defect in pyruvate metabolism, rather than a global defect in central carbon metabolism (**Figure 3B**). Similarly, *FBXW7*-depleted HEK293T spheroids failed to grow in glutamine-free conditions, even after pyruvate supplementation (**Figure 3C, D**). To clarify the fate of pyruvate in *FBXW7*-depleted cells, we repeated our ^13^C pyruvate tracer experiment and focused on malate and aspartate, whose relative fractions of m+3 isotopologue increased in glutamine-limited conditions (**Figure 1L**). Importantly, while we confirmed increased enrichment in m+3 isotopologue (following direct conversion of ^13^C pyruvate to oxaloacetate) in the absence of glutamine, we found a significant reduction in the same pool in *FBXW7*-depleted cells, irrespective of the presence of glutamine, further suggestive of impaired pyruvate metabolism (**Figure 3E, F; Table S6**).

As the primary function of FBXW7 is known to be in protein degradation, we performed a global cell proteomics analysis to understand how this tumor suppressor and glutamine deprivation affect the proteome. We observed only a few changes in either condition, but importantly, we found that PC was among the most significantly decreased proteins in *FBXW7*-depleted cells, again independently of the presence of glutamine, and common to multiple cell lines (**Figure 3G, H; Figure S2; Tables S7-9**). *PC* was one of the best hits of the screen (**Figure 3A**), and low PC expression is fully compatible with both the defects in pyruvate metabolism and the impaired growth in the absence of glutamine seen in cells lacking *FBXW7*. To test for a causal role, we expressed a *PC* cDNA in *FBXW7*-depleted cells. We found that *PC* overexpression not only conferred a growth advantage to cells during glutamine deprivation, it also rescued the proliferation of *FBXW7*-depleted cells to the same extent as in control cells, suggestive of an epistatic effect (**Figure 3H, I**). Taking these results together, we propose that *FBXW7* supports the proliferation of cells in the absence of glutamine by promoting PC expression and subsequently pyruvate anaplerosis.

### An FBXW7-MYC-MNT-SIN3A axis controls the expression of PC via histone deacetylation

Our observation that PC protein abundance is reduced in the absence of *FBXW7* was unanticipated, firstly because FBXW7 is considered to be mainly involved in protein degradation, and secondly because the two proteins are not localized in the same cellular compartment^23–25^. To investigate this further, we measured the transcript levels of *PC* and observed that they were significantly decreased in *FBXW7*-depleted cells (**Figure 4A**), mirroring the effect observed in immunoblotting experiments (**Figure 3H**). Intriguingly, PC mRNA stability was not affected, suggesting a decrease in its rate of transcription (**Figure 4B**).

**Figure 4.**
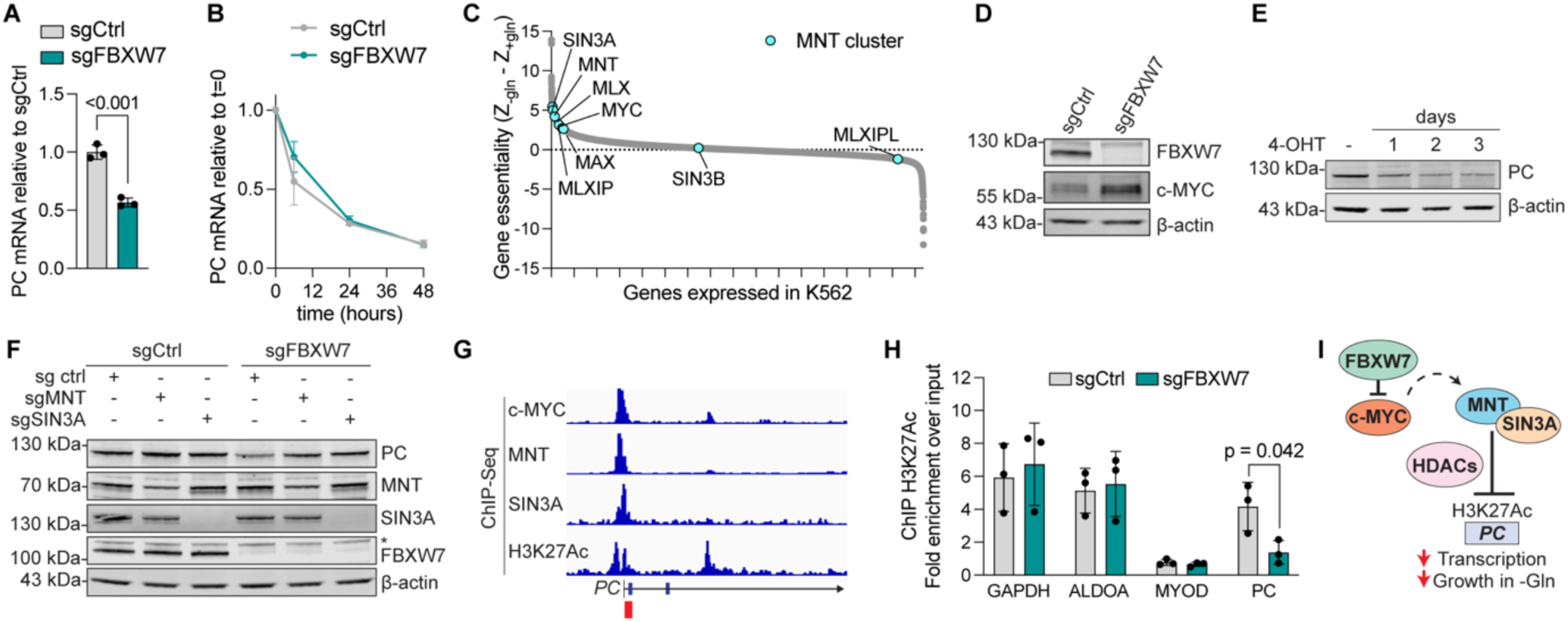
PC expression is repressed by c-MYC and MNT-/SIN3A-dependent histone deacetylation. **(A)** qPCR for PC mRNA expression in control and *FBXW7*-depleted cells. p value: unpaired t-test. **(B)** mRNA decay in actinomycin-treated samples. **(C)** c-MYC and the MNT cluster are hits in the glutamine-deprivation CRISPR/Cas9 screen. (**D)** Immunoblot displaying c-MYC expression in control and *FBXW7*-depleted cells. **(E)** Immunoblot showing reduced PC expression in MycER mouse embryonic fibroblasts treated with 4-hydroxytamoxifen (4-OHT) over the course of 3 days. **(F)** Immunoblot showing PC expression in *MNT* or *SIN3A*-depleted sgCtrl and sgFBXW7 cells. *=aspecific band. **(G)** ChIP-seq data (source: ENCODE) showing c-MYC, MAX, MNT and SIN3A predicted binding sites as well as H3K27Ac peaks on the *PC* promoter. Red: region amplified by our qPCR assay used for validation. **(H)** ChIP-qPCR for H3K27Ac on *GAPDH*, *ALDOA*, *MYOD* and *PC* promoters. p values: multiple paired t-tests. **(I)** Proposed model of *PC* transcriptional regulation.

We reasoned that additional factors involved in *PC* expression might also affect proliferation in the absence of glutamine, and we mined our genome-wide CRISPR/Cas9 screen to identify transcription-related factors. We found a cluster of epigenetic modifiers whose depletion is beneficial for growth in the absence of glutamine (**Figure 4C**) that included *MNT* and its binding partners *MAX*, *MLX* and *SIN3A*, as well as *MLXIP*, all of which are involved in transcriptional repression^26^. Notably, this “MNT cluster” lies downstream of c-MYC, an FBXW7 target that can skew the proportion of MAX, MNT and MLX dimers^19,26^, and we confirmed that c-MYC accumulates in *FBXW7*-depleted K562 cells (**Figure 4D**). As *MYC* is in the top 1% essential genes in K562^27^, we first focused on the effects of its overexpression, rather than depletion. To this aim, we employed the tamoxifen-inducible MycER mouse embryonic fibroblast line, which has extensively been used to study *MYC*-dependent glutamine dependency^7,13^. Importantly, we found that *MYC* overexpression reduced PC protein abundance (**Figure 4E**), mirroring the results obtained in *FBXW7*-depleted K562. Furthermore, depletion of *MNT* or *SIN3A* (acting downstream of c-MYC) was sufficient to rescue expression of PC in a *FBXW7*-depleted background (**Figure 4F**), thus confirming regulation of *PC* by MYC and the MNT cluster.

Genes from the MNT cluster act as transcriptional repressors that bind genomic DNA and recruit histone deacetylases (HDACs) to lysine 27 of histone 3 (H3K27)^28,29^. This prompted us to probe for H3K27 acetylation, one of the main epigenetic alterations in cancers driven by *FBXW7* loss of function^30^. We first analyzed the chromatin immunoprecipitation (ChIP) and sequencing data from the ENCODE project^31^ and observed a clear signature for binding of c-MYC, MNT and SIN3A on the *PC* promoter in K562 cells, as well as H3K27Ac peaks (**Figure 4G**). We next performed our own ChIP experiment using a set of qPCR primers designed specifically to measure H3K27Ac on the *PC* promoter (**Figure 4G**, red bar). We found that while the H3K27Ac profile of control genes with either high (*GAPDH*, *ALDOA*) or low (*MYOD*) acetylation was unaffected, acetylation on the *PC* promoter was significantly reduced in *FBXW7*-depleted cells (**Figure 4H**), again mirroring transcript and protein abundance. Taken together, our data indicate that upon increased c-MYC expression in *FBXW7*-depleted cells, an MNT-related cluster of transcriptional repressors recruits HDACs on the *PC* promoter, leading to decreased H3K27 acetylation, low *PC* expression, impaired pyruvate anaplerosis, and glutamine dependency (**Figure 4I**).

## Discussion

Glutamine is one of the most versatile and highly consumed amino acids. Here, we have employed parallel nutrient and genetic screening to provide a comprehensive inventory of metabolites and molecular pathways required for the proliferation of cells in glutamine-deprived environments. Previous efforts had identified nutrients that could partially compensate for glutamine depletion, or glutaminase inhibition, including carboxylic acids, aspartate and asparagine^10–12,14^. Our systematic approach has confirmed the role of some of these metabolites, and in addition provides a direct comparison between hundreds of nutrients, covering most of the diversity found in the plasma or in the tumor interstitial fluid^2,32^. Our data highlight pyruvate and TCA cycle-related metabolites as the main determinants for cell proliferation in absence of glutamine. While glutamine is considered both a carbon and a nitrogen source, our findings indicate that its involvement in central carbon metabolism outweighs its role as a nitrogen donor, possibly due to the presence of other nitrogen-containing molecules, such as amino acids, in the environment. We confirmed the role of pyruvate in carbon metabolism using a ^13^C-labeled tracer, which showed significant incorporation of pyruvate-derived carbon in the TCA cycle intermediates as well as in aspartate and asparagine, two amino acids essential for proliferation. In agreement with earlier studies, we could show that supplementation with aspartate or asparagine was sufficient to restore proliferation in glutamine-restricted cells^11,12,14^.

To extend this analysis we took advantage of the observation that pyruvate-supplemented cells are able to survive glutamine starvation to design a genome-wide CRISPR/Cas9 screen aimed at identifying the genes involved in the proliferation of glutamine-restricted cells in the presence of pyruvate. Integrating data from our nutrient and genetic screens with our ^13^C pyruvate tracer analysis and earlier literature, we can now propose a unified model in which pyruvate rescues cell proliferation in glutamine-limiting conditions by serving as an anaplerotic substrate for PC, thus promoting mitochondrial aspartate biogenesis, its subsequent export to the cytosol via the malate-aspartate shuttle, and its utilization in asparagine, protein and nucleotide synthesis (**Figure 2J**). In support of this model, we showed that while depleting the mitochondrial arm of the malate-aspartate shuttle, involved in aspartate export, is detrimental in glutamine-limited conditions, depleting its cytosolic arm, involved in bringing carbon back into mitochondria, is beneficial. Similarly, we report that inhibiting the plasma membrane cystine/glutamate exchanger system X_c_^-^ is beneficial to cell growth when glutamine is limiting. This observation could be explained by conservation of intracellular glutamine-derived glutamate and warrants caution in the use of system X_c_^-^ inhibitors that are currently being developed for cancer therapy, as they could promote the survival of cells in the glutamine-limited tumor microenvironment. Interestingly, our work also highlights the need for the vitamin biotin, and the machinery for biotin import and conjugation to proteins, which provides an essential co-factor for PC. Together, our findings extend previous work, and in addition provide a comprehensive resource focusing on both the metabolites and the genes which mediate survival of glutamine-starved cells.

Our analysis also highlighted the tumor suppressor gene *FBXW7* as a central regulator of pyruvate metabolism. *FBXW7* is one of the most highly mutated genes in bile duct, blood, endometrium, colon and stomach cancer^33,34^, and as an E3 ubiquitin ligase, it degrades the protein product of several proto-oncogenes including c-MYC, c-JUN and cyclin E^19–22^. *FBXW7* also influences epigenetic remodeling via H3K27Ac and H3K27me3 by acting upstream of MYC^30^. *FBXW7* has been linked to oxidative and liver metabolism^35,36^, but the mechanism behind these observations remained unclear. Here, we report that *FBXW7* mediates anaplerosis by promoting the expression of the mitochondrial pyruvate carboxylase *PC*. *FBXW7*-depleted cells show glutamine addiction that cannot be rescued by pyruvate supplementation, thus leading to decreased carbon flux to sustain aspartate and asparagine synthesis. We confirmed that c-MYC accumulates in *FBXW7*-depleted cells, and we report here that *PC* transcription decreases in parallel. c-MYC acts upstream of MNT and SIN3A, two epigenetic repressors that bind the promoter of *PC* and that are known to recruit histone deacetylases to chromatin^28,29^. Here, we showed decreased H3K27 acetylation of the *PC* promoter, lower *PC* expression and reduced pyruvate utilization in the TCA cycle, thus limiting metabolic flexibility and preventing survival upon glutamine starvation in *FBXW7*-depleted cells. c-MYC is a known driver of glutamine addiction that promotes expression of glutaminase and glutamine transporters^6–8^. Despite MYC accumulation and glutamine dependency of K562 cells, our proteomics analysis of *FBXW7*-depleted cells did not identify changes in the expression of these glutamine-related genes, suggesting cell-specific differences (**Figure 3G; Tables S8, S9**). Rather, we observed reduced PC levels, and importantly showed that *PC* overexpression restores growth of *FBXW7*-depleted cells, indicating that *PC* is both necessary and sufficient to promote proliferation upon glutamine depletion. Our findings provide a better understanding of metabolic pathways regulated by *FBXW7* and highlight PC as a novel effector downstream of FBXW7/c-MYC involved in pyruvate carboxylation and glutamine addiction, two related metabolic defects that could be targeted for therapy.

Altogether, our study highlights the power of nutrient-based screening coupled with genetic perturbation strategies to provide a better understanding of the metabolic flexibility of mammalian cells facing nutrient deprivation. Our results provide a unified model of the molecular mechanisms which allow proliferation of glutamine-starved cells, as well as an important resource for further investigations in the field. Glutamine depletion, and nutrient fluctuations in general, are constantly faced by mammalian cells according to both diet and changing physiological states, such as tissue migration, cell growth and differentiation. Investigating how cells cope with such heterogenous conditions will increase our understanding of the factors supporting metabolic flexibility, and could provide applications in pathological settings such as nutrient scarcity upon pathogen infection and malignant cell adaptation to the cancer microenvironment.

## Limitations of the study

The present work provides a systematic analysis of both nutrients and molecular pathways required to sustain proliferation of cells in a glutamine-restricted environment. A common limitation of genetic screens performed in a pooled format is cross-feeding (namely the transfer of nutrients between cells via the culture medium) and competition between cells in the population. While we have validated many of the hits from our screens, future work should include continuing the experimental validation of the additional genes and metabolites highlighted by these approaches. Our work thus far has focused on seven adherent and suspension cancer cell lines grown in two or three dimensions. In the future, it will be important to validate our findings in primary cells and in cancer models. Our findings also assigned a role for *FBXW7* in *PC* regulation and pyruvate metabolism, and while we focused on c-MYC and its well-characterized MNT/SIN3A effectors^26^, both FBXW7 and c-MYC have a wide range of targets. We do not exclude that additional layers of regulation may exist downstream of *FBXW7* and upstream of *PC* that could contribute to pyruvate anaplerosis and cell proliferation in glutamine-deprived conditions.

## Acknowledgements

The authors would like to thank past and present members of the Jourdain laboratory, M. Knobloch, F. Langlet, J.C. Martinou, K. Maundrell, A. Mayer, L. Mazzeo, M. Quadroni, O.S. Skinner, and the Metabolomics Platform and the Protein Analysis Facility (University of Lausanne). MycER mouse embryonic fibroblasts were a kind gift from Raul Mostoslavsky (Harvard Medical School). This project was supported by an EMBO Postdoctoral fellowship (ALTF286-2022, to M.L.) and the Swiss National Science Foundation (310030_200796, to A.A.J.).

## Author contributions

**ML:** investigation (lead); methodology (equal); validation (lead); data curation (lead); project administration (equal); formal analysis (lead); visualization; funding acquisition (supporting); writing (original draft preparation, equal). **FV:** investigation (supporting); data curation (supporting); writing (reviewing and editing, equal). **HGA:** formal analysis (supporting); data curation (supporting). **JI:** methodology (equal); formal analysis (supporting); data curation (supporting); writing (reviewing and editing, equal). **AAJ:** conceptualization; methodology (equal); project administration (equal); supervision (lead); funding acquisition (lead); writing (original draft preparation, equal).

## Declaration of generative AI and AI-assisted technologies

No generative AI or AI-assisted technology was employed for conceptualization or investigation in the course of this project.

## Supplemental information

**Figure S1.**
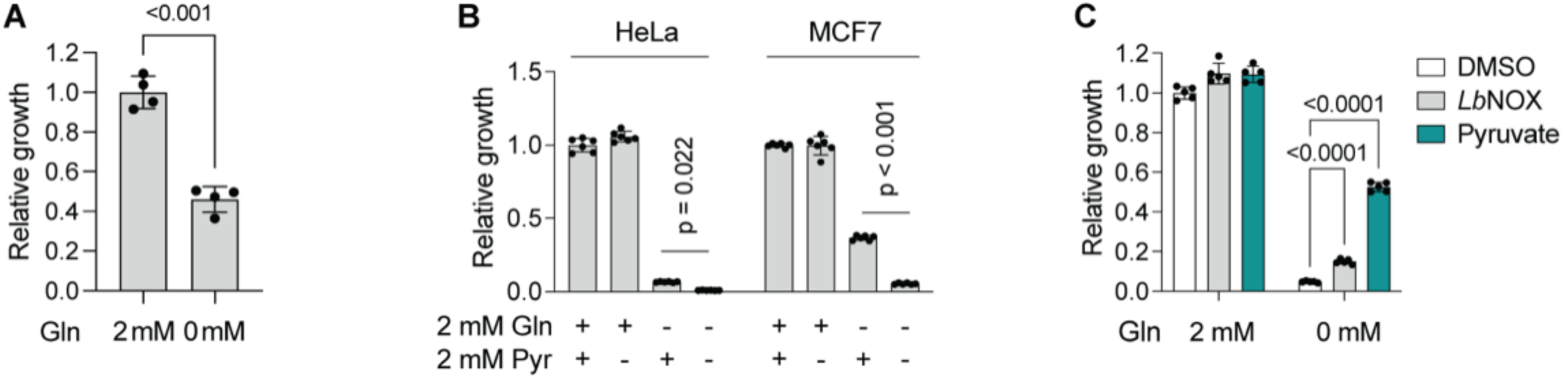
Proliferation of different cell lines in glutamine-deprived conditions upon pyruvate supplementation or *Lb*NOX overexpression. **(A)** CellTiter-Glo growth assay of K562 cells in medium with or without glutamine (Gln). p values: unpaired t-test. **(B)** Proliferation of HeLa and MCF7 cells cultured in medium supplemented with (+) or depleted of (-) glutamine or pyruvate. p values: one-way ANOVA. **(C)** Growth of K562 cultured in medium supplemented with or deprived of glutamine. Cells were either overexpressing *Lb*NOX or cultured in medium supplemented with pyruvate. p values: one-way ANOVA.

**Figure S2.**
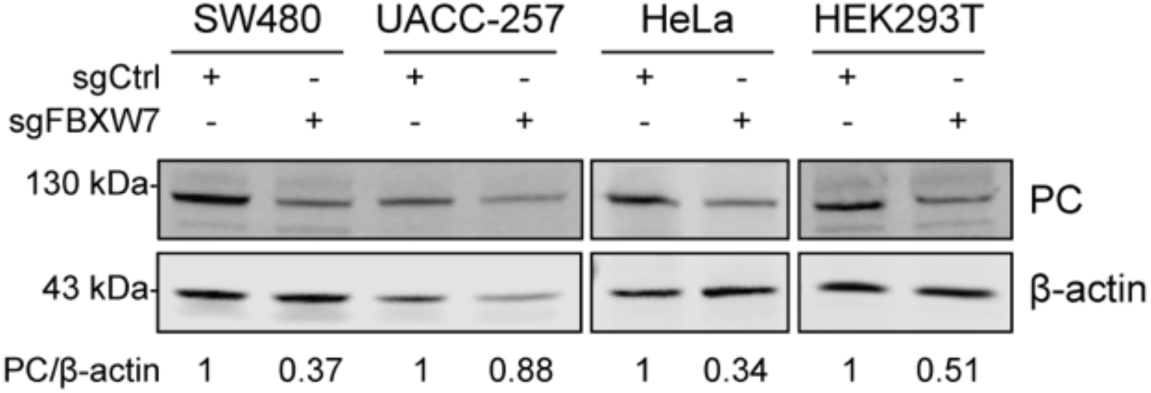
Loss of *FBXW7* leads to PC downregulation in multiple cell lines. Immunoblot showing reduced PC expression in *FBXW7*-depleted SW480, UACC-257, HeLa and HEK293T cells. Relative quantification of PC abundance (fluorescent antibody signal, normalized to β-actin) is displayed below each lane.

Table S1. Results of nutrient-sensitized screen in glutamine-deprived media.

Table S2. Multiple pathways metabolomics analysis under glutamine starvation.

Table S3. Multiple pathways metabolomics analysis in glutamine-deprived media enriched with pyruvate.

Table S4. ^13^C pyruvate isotope tracing in glutamine-rich and glutamine-deprived medium.

Table S5. Results of CRISPR/Cas9 screen in glutamine-deprived media.

Table S6. ^13^C pyruvate isotope tracing in *FBXW7*-depleted cells in glutamine-rich and glutamine-deprived medium.

Table S7. Protein mass spectrometry of sgCtrl cells in glutamine-rich and glutamine-deprived conditions.

Table S8. Protein mass spectrometry of *FBXW7*-depleted cells in glutamine-rich medium.

Table S9. Protein mass spectrometry of *FBXW7*-depleted cells in glutamine-deprived medium.

Table S10. List of sgRNAs and sequences used in this study.

Table S11. List of primers used for ChIP-qPCR.

## Methods

### Experimental models

K562 (ATCC, CCL-243), HEK293T (ATCC, CRL-3216), HeLa (ATCC, CCL-2), MCF7 (ATCC, HTB-22), SW480 (ATCC, CCL-228) and UACC-257 (DCTD, CVCL_1779) were maintained under standard cell culture conditions in DMEM containing 25 mM glucose, 1 mM sodium pyruvate, 2 mM glutamine and supplemented with 10% fetal bovine serum (FBS) and 100 U/mL penicillin/streptomycin. MycER mouse embryonic fibroblasts were a kind gift from Raul Mostoslavsky (Harvard Medical School) and were cultured in the same medium as indicated above. *Myc* expression was induced by supplementing the cell culture medium with 200 nM 4-hydroxytamoxifen (4-OHT) (MedChemExpress, HY-16950). All cells were cultured at 37°C, 5% CO_2_ and regularly tested to ensure absence of mycoplasma.

### Spheroid culture

HEK293T spheroids were seeded in DMEM supplemented with 25 mM glucose, 10% dialyzed FBS and 100 U/mL penicillin/streptomycin. Experimental media were supplemented with either 2 mM pyruvate or 2 mM glutamine, as indicated. HEK293T were seeded on U-bottom Nunclon^TM^ Sphera^TM^ 96-Well plates (ThermoFisher, 174925) and briefly centrifuged to allow cells to settle at the bottom of the well. Samples were imaged with an EVOS^TM^ FL imaging system (Life Technologies) every 3-4 days with a 4x objective. Spheroid diameter was quantified using Fiji (ImageJ). For biotin and pyruvate supplementation, HEK293T cells were first cultured in standard DMEM (25 mM glucose, 1 mM pyruvate, 2 mM glutamine) supplemented with 100 U/mL penicillin/streptomycin and 10% dialyzed FBS for two weeks to deprive cells of biotin. After 14 days, HEK293T were seeded in DMEM containing 25 mM glucose and supplemented with 10% dialyzed FBS and 100 U/mL penicillin/streptomycin. Test media were prepared with the addition of either 2 mM of pyruvate, 1 µM of biotin, or both, as indicated in the figure legends.

### Proliferation assays

Cells were seeded at 0.1 million/ml in DMEM supplemented with 25 mM glucose, 10% dialyzed FBS and 100 U/mL penicillin/streptomycin in black Nunc^TM^ MicroWell^TM^ 96-Well plates (NUNC, 137101) for the Prestoblue assay, or in flat-bottom 12-well plates for standard cell and viability count. Test media were supplemented with 2 mM glutamine, 2 mM pyruvate, or 2 mM of all other amino acids or carboxylic acid as indicated, with the exception of aspartate and asparagine, which were added at a final concentration of 150 µM as previously described^37^. Proliferation was measured after 4-5 days either by trypan-blue mediated count (Vi-cell blu counter, Beckman Coulter), or by adding the Prestoblue dye (Life Technologies, A13262) and measuring fluorescence (excitation: 560 nm, emission: 590 nm) or by using the CellTiter-Glo dye (Promega, G9242) and measuring luminescence, with the last two performed using a BioTek Synergy plate reader (Agilent). “Relative growth” was quantified by normalizing proliferation to the internal control sample (complete media, untreated cells, or sgCtrl). For drug treatments, 375 µM sulfasalazine or 100 nM ISRIB were added to the cell culture medium at the beginning of the growth assay as indicated.

### Cloning and lentiviral infection

Gene-specific guides (sgRNA) were selected from the two best sequences as scored in the CRISPR/Cas9 screen and were ordered as complementary oligonucleotides (Integrated DNA Technologies, IDT). sgRNAs were cloned into the pLentiCRISPR v2 vector (Addgene, 52961) using the BsmBI site. Two guides targeting olfactory receptors not expressed in K562, *OR2M4* and *OR11A1*, were used to generate vectors for the negative control (sgCtrl) samples. Plasmids pLX304-GFP-V5 (Addgene, 193687) and pLX304-PC (DNAsu, HsCD00436386) were used for gene overexpression. Lentiviruses were produced in HEK293T cultures, as previously described^38^ and K562 cells were infected in the presence of 10 µg/ml polybrene (Sigma-Aldrich, TR-1003G). Successfully infected K562 were selected after 24 h using 2 µg/ml puromycin (pLentiCRISPR v2) or 100 µg/mL blasticidin (pLX304). Cells were counted 3 days post-selection and maintained in standard cell culture medium as described above for 10-12 days before analysis to ensure gene knockout and target protein degradation, or 7 days for overexpression, both of which were confirmed by immunoblotting. sgRNA and primer sequences are described in Table S10.

### Genome-wide CRISPR/Cas9 screen

K562 were infected with the lentiviral Brunello library (Genome Perturbation Platform, Broad Institute) containing 76,441 sgRNAs^39^ in duplicates, as previously described^40^. Infection was performed at multiplicity of infection 0.3, using 500 cells per sgRNA guide, and in the presence of 10 µg/ml polybrene (Sigma-Aldrich, TR-1003G). Successfully infected cells were selected after 24 h using 2 µg/ml puromycin. After 10 days, cells were plated at 0.1 million/mL (equivalent to 1000 cells per sgRNA for the glutamine-rich condition, and 2000 cells per sgRNA for the glutamine-depleted condition) in DMEM medium supplemented with 10% dialyzed FBS, 100 U/mL penicillin/streptomycin, 25 mM glucose, 50 µg/ml uridine and either 2 mM glutamine or water as a negative control. Cells were cultured for 3 weeks and harvested on day 31 post-infection. The NucleoSpin Blood kit (Machery Nagel, 740954.20) was used to extract total genomic DNA, which was subsequently used for barcode sequencing, mapping and read count at the Genome Perturbation Platform (Broad Institute).

Screen results were analyzed as previously outlined^18^ using normalized Z scores. In brief, raw sgRNA read counts were normalized to reads/million and log_2_ transformed. The mean log_2_(fold change) was calculated based on read counts in the pre-swap control (day 7 post-infection) and averaging the abundance of four sgRNAs. Results were averaged based on the mean across two replicates. K562 RNA-sequencing data (sample GSM854403 in Gene Expression Omnibus series GSE34740) were used to to filter out genes not expressed in K562. Z score was calculated from the mean log_2_(fold change) using the statistics of the null distribution, defined by the 3726 genes with lowest expression.

### BioLog nutrient screen

Cells were seeded at 0.1 million/mL in the PM-M1, PM-M2, PM-M3 and PM-M4 BioLog plates (Biolog, PM-M1: 13101, PM-M2: 13102, PM-M3: 13103, PM-M4: 13104) in DMEM containing 25 mM glucose and supplemented with 10% dialyzed FBS and 100 U/mL penicillin/streptomycin. Samples were cultured for 4 days at 37°C, 5% CO_2_. Cell growth was measured using the BioLog MA dye (Biolog, 74351) by quantifying absorbance at 590 nm using a SpectraMax ID3 plate reader (Molecular Devices).

### Immunoblotting

Cells were washed in ice-cold PBS and flash-frozen on dry ice as pellets. Samples were lysed in RIPA buffer (25 mM Tris pH 7.5, 150 mM NaCl, 0.1% SDS, 0.1% sodium deoxycholate, 1% NP40 analog) supplemented with cOmplete^TM^ ULTRA protease inhibitor tablets (Roche, 05892970001) and Pierce^TM^ Universal Nuclease (Life Technologies, 88702) by incubation on ice for 20 min. Supernatants were cleared by spinning samples for 10 min at 14,000 *g* at 4°C. Protein concentration was quantified using DC Protein Assay (Bio-rad). Lysates were loaded on 8-16% Novex^TM^ WedgeWell^TM^ Tris-Glycine gels (ThermoFisher, XP08160BOX) and run at 200 V for 1 h. Samples were transferred to nitrocellulose membranes using a wet transfer chamber overnight at 4°C (60 V). Membranes were blocked in 5% milk diluted in TBS + 0.1% Tween (TBS-T) and incubated with a 1:500-1:1000 dilution of primary antibody for 1.5 h at room temperature (or overnight at 4°C), then with a secondary antibody at 1:10,000 dilution for 1 h at room temperature. All antibodies were diluted in 5% milk in TBS-T. Washes were performed in TBS-T and membranes were imaged using fluorescence detection on an Odyssey CLx analyzer (Li-Cor).

### RNA extraction, cDNA synthesis and qPCR

Total RNA was extracted in phenol/chloroform using TRI reagent (Sigma-Aldrich, T3809), as described by the manufacturer. Complementary DNA (cDNA) was synthesized using random primers and M-MLV Reverse Transcriptase (Sigma-Aldrich, M1302-40KU) in the presence of RNAseOUT Recombinant RNAse inhibitor (Invitrogen, 10777019) and random primers (ThermoFisher, N8080127). Quantitative PCR (qPCR) was performed using Taqman probes (see STAR Methods table) and a CFx96 quantitative PCR machine (Bio-Rad). All data were internally normalized to *TBP* expression using the ΔΔCt method.

For mRNA decay analysis, cells were treated with 1 µg/ml actinomycin D (Sigma-Aldrich, A9415) and harvested at the indicated time points. Total RNA extraction, cDNA synthesis and qPCR were performed as described above. Relative mRNA decay was quantified by normalizing ΔΔCt values at each time point to the initial ΔΔCt (t=0, untreated cells). Plot shows data points from two independent replicates.

### Chromatin immunoprecipitation (ChIP)-qPCR

For chromatin isolation, 10 million cells were harvested per sample, washed in PBS and fixed in 1% formaldehyde for 10 min at room temperature. Samples were quenched using 125 mM glycine, washed in PBS, lysed in ChIP Lysis buffer (50 mM Hepes-KOH pH 7.5, 140 mM NaCl, 1 mM EDTA pH 8, 1% Triton X-100, 0.1% sodium deoxycholate and 0.1% SDS) and sonicated using a Bioruptor (Diagenode) to obtain 100-300 bp fragments. Lysates were cleared by centrifugation at maximum speed for 10 min at 4°C, and an aliquot from each was retained as the input sample.

For chromatin immunoprecipitation, lysates were incubated overnight at 4°C with 5 µg IgG control (Abcam, ab171870) or α-H3K27Ac (Abcam, ab4729). The next day, samples were incubated with protein G beads (previously blocked in 10% bovine serum) for 2 h at room temperature. Beads were washed using wash buffer (20 mM Tris-HCl pH 8, 150 mM NaCl, 2 mM EDTA pH 8, 1% Triton X-100, 0.1% SDS). The final wash was performed with a higher salt concentration (500 mM NaCl) to improve the signal to noise ratio. After elution, samples were treated with RNAse A (Takara Bio, 740505) for 1 h at 37°C and proteinase K (Sigma-Aldrich, SRE0005) first for 1 h at 55°C, then overnight at 65°C to remove contaminants. DNA cleanup was performed using phenol-chloroform-isoamyl alcohol (Sigma-Aldrich, 77617) as described by the manufacturer.

For qPCR, primers were designed using the NCBI genome data viewer (https://www.ncbi.nlm.nih.gov/gdv/) and the Integrative Genomic Viewer (IGV) software, GRCh38.p14 Primary Assembly. Oligonucleotides were obtained via Integrated DNA Technologies (IDT). qPCR was performed using primers for the promoter of the *PC* gene, as well as primers for genes showing low (*MYOD*) or high (*GAPDH*, *ALDOA*) levels of H3K27Ac in K562 cells as controls. Data was normalized to the signal in the input sample using the ΔΔCt method. A list of primers used for ChIP-qPCR is provided in Table S11.

### Protein mass spectrometry

Protein mass spectrometry was performed at the Protein Analysis Facility at the University of Lausanne. Samples were lysed in 250 µL miST lysis buffer (1% Sodium deoxycholate, 100 mM Tris pH 8.6, 10 mM DTT), heated for 10 min at 75°C and digested following a modified version of the iST method^41^. Based on tryptophan fluorescence quantification^42^, 100 µg of proteins at 2 µg/µl in miST buffer were heated 5 min at 95°C, diluted 1:1 (v:v) with water containing 4 mM MgCl_2_ and 250 Units/µL benzonase (Merck, 70746) and incubated for 15 min at room temperature to digest nucleic acids. Reduced disulfides were alkylated by adding one fourth of the vol. of 160 mM chloroacetamide (32 mM final) and incubating for 45 min at room temperature in the dark. Samples were adjusted to 3 mM EDTA and digested with 1 µg Trypsin/LysC mix (Promega, V5073) for 1 h at 37°C, then for 1 h with an additional 0.5 µg of proteases. To remove sodium deoxycholate, two sample volumes of isopropanol containing 1% TFA were added to the digests, and the samples were desalted on a strong cation exchange (SCX) plate (Oasis MCX; Waters Corp., Milford, MA) by centrifugation. After washing with isopropanol/1%TFA, peptides were eluted in 200 µl of 80% MeCN, 19% water, 1% (v/v) ammonia, and dried by centrifugal evaporation.

Aliquots (1/8) of each sample were pooled and separated into 6 fractions by off-line basic reversed-phase (bRP) using the Pierce High pH Reversed-Phase Peptide Fractionation Kit (ThermoFisher). The fractions were collected in 7.5, 10, 12.5, 15, 17.5 and 50% acetonitrile in 0.1% triethylamine (∼pH 10). Dried bRP fractions were redissolved in 50 ul 2% acetonitrile with 0.5% TFA, and 5 µl were injected for LC-MS/MS analyses. LC-MS/MS analyses were carried out on a TIMS-TOF Pro (Bruker, Bremen, Germany) mass spectrometer interfaced through a nanospray ion source (“captive spray”) to an Ultimate 3000 RSLCnano HPLC system (Dionex). Peptides were separated on a reversed-phase custom packed 45 cm C18 column (75 μm ID, 100Å, Reprosil Pur 1.9 µm particles, Dr. Maisch, Germany) at a flow rate of 0.250 µl/min with a 2-27% acetonitrile gradient in 93 min followed by a ramp to 45% in 15 min and to 90% in 5 min (total method time: 140 min, all solvents contained 0.1% formic acid). Identical LC gradients were used for DDA and DIA measurements.

For creation of the spectral library, data-dependent acquisitions (DDA) were carried out on the 6 bRP fractions sample pool using a standard TIMS PASEF method^43^ with ion accumulation for 100 ms for each survey MS1 scan and the TIMS-coupled MS2 scans. Duty cycle was kept at 100%. Up to 10 precursors were targeted per TIMS scan. Precursor isolation was done with a 2 Th or 3 Th window below or above m/z 800, respectively. The minimum threshold intensity for precursor selection was 2500. If the inclusion list allowed it, precursors were targeted more than one time to reach a minimum target total intensity of 20’000. Collision energy was ramped linearly based uniquely on the 1/k0 values from 20eV (at 1/k0=0.6) to 59 eV (at 1/k0=1.6). Total duration of a scan cycle including one survey and 10 MS2 TIMS scans was 1.16 s. Precursors could be targeted again in subsequent cycles if their signal increased by a factor 4.0 or more. After selection in one cycle, precursors were excluded from further selection for 60 s. Mass resolution in all MS measurements was approximately 35’000. The data-independent acquisition (DIA) used essentially the same instrument parameters as the DDA method reported previously^44^. Per cycle, the mass range 400-1200 m/z was covered by a total of 32 windows, each 25 Th wide and a 1/k0 range of 0.3. Collision energy and resolution settings were the same as in the DDA method. Two windows were acquired per TIMS scan (100 ms) so that the total cycle time was 1.7 s.

Raw Bruker MS data were processed with Spectronaut 16.2 (Biognosys, Schlieren, Switzerland). A library was constructed from the DDA bRP fraction data by searching the annotated human proteome SWISSPROT database of January 7th, 2022 (20’375 sequences). For identification, peptides of 7-52 AA length were considered, cleaved with trypsin/P and a maximum of 2 missed cleavages were allowed. Carbamidomethylation of cysteine (fixed), methionine oxidation and N-terminal protein acetylation (variable) were the modifications applied. Mass calibration was dynamic and based on a first database search. The Pulsar engine was used for peptide identification. Protein inference was performed with the IDPicker algorithm. Spectra, peptide and protein identifications were all filtered at 1% false discovery rate (FDR) against a decoy database. Specific filtering for library construction removed fragments corresponding to less than 3 AA and fragments outside the 300-1800 m/z range. Only fragments with a minimum base peak intensity of 5% were kept. Precursors with less than 3 fragments were also eliminated and only the best 6 fragments were kept per precursor. No filtering was done on the basis of charge state and a maximum of 2 missed cleavages was allowed. Shared (non proteotypic) peptides were kept. The library created contained 110’276 precursors mapping to 83’238 stripped sequences, of which 79’395 were proteotypic. These corresponded to 7’502 protein groups (7’613 proteins). Of these, 806 were single hits (one peptide precursor). In total, 649’960 fragments were used for quantitation.

Peptide-centric analysis of DIA data was done with Spectronaut 16.2 using the library described above. Single-hit proteins (defined as being matched by one stripped sequence only) were kept in the Spectronaut analysis. Peptide quantitation was based on XIC area, for which a minimum of 1 and a maximum of 3 (the 3 best) precursors were considered for each peptide, from which the median value was selected. Quantities for protein groups were derived from inter-run peptide ratios based on the MaxLFQ algorithm^45^. Global normalization of runs/samples was done based on the median of peptides. Overall, 105’024 precursors were quantified in the dataset, mapping to 7’190 protein groups. 96’612 precursors (6’889 protein groups) had full profiles, i.e. were quantified in all samples. The average number of data points per peak was 7.4.

Subsequent analyses were done with the Perseus software package (version 1.6.15.0)^46^. Contaminant proteins were removed, and quantity values were log_2_-transformed. After assignment to groups, only proteins quantified in at least 3 samples of the single-hit group were kept. Volcano plots were generated on Prism (GraphPad Software) after Benjamini-Hochberg false discovery rate (FDR) correction. All raw MS data together with raw output tables are available via the Proteomexchange data repository (www.proteomexchange.org).

### Multiple pathway metabolomics analysis of polar metabolites

Metabolomics analysis was performed at the Metabolomics Platform at the University of Lausanne. Cell pellets were extracted with 80% methanol, sonicated and homogenized (Precellys Cryolys). Lysates were centrifuged at 15000 rpm at 4°C for 15 min, evaporated to dryness and reconstituted in methanol, based on total protein content (quantified by prior BCA assay). Samples were analyzed by liquid chromatography coupled to mass spectrometry (LC-MS/MS) following previously described methods^47^.

For metabolite relative quantification, ultra-high performance liquid chromatography coupled to tandem mass spectrometry (UHPLC-MS/MS) was used, and a Triple Quadrupole mass spectrometer (6495 iFunnel Agilent) in multiple reaction monitoring (MRM) mode was employed for targeted measurement, as previously detailed^47,48^. Metabolome coverage was maximized by using two liquid chromatography modes coupled to positive and negative electrospray ionization MS^49^.

Raw LC-MS/MS data was analyzed using Agilent Quantitative analysis software version B.07.00 (MassHunter Agilent technologies). Extracted Ion Chromatogram (EIC) areas were used for monitored MRM transitions for relative metabolite quantification. All tables containing peak areas of detected metabolites were processed and filtered based on CV calculated across QC samples. Data was discarded when peaks showed analytical variability with CV above 30%. False discovery rate (FDR) correction was performed using the Benjamini, Krieger and Yekutieli method. Metabolites with FDR<1% were loaded on MetaboAnalyst 6.0 for Pathway Impact Analysis using the Homo sapiens KEGG pathway library^50^. Both metabolite abundance and Pathway Impact Analysis results were plotted using Prism (GraphPad Software).

### ^13^C pyruvate isotopic profiling

^13^C pyruvate isotopic profiling was performed at the Metabolomics Platform at the University of Lausanne. Cell extracts were prepared and quantified as above (see “Multiple pathway targeted analysis of polar metabolites”) and analyzed using Hydrophilic Interaction Liquid Chromatography coupled to high resolution mass spectrometry (HILIC-HRMS) in negative ionization mode with a 6550 Quadrupole Time-of-Flight (Q-TOF) system interfaced with 1290 UHPLC system (Agilent Technology), as detailed elsewhere^49^. Metabolite separation was achieved via an iHILIC-(P) Classic PEEK column (100 x 2.1 mm, 5 μm particle size, HILICON AB, Umea, Sweden) column. Mobile phase composition was A = 20 mM ammonium acetate and 20 mM NH4OH in water at pH 9.7 and B = 100% ACN. Linear gradient elution: 90% B (0-1.5 min) to 50% B (8-11 min) down to 45% B (12-15 min). For column re-equilibration, the initial chromatographic conditions were set during 9 min post-run (flow rate 300 μL/min, column temperature 30°C, injection volume 2 µl). ESI conditions were the following: dry gas temperature 290°C and flow 14 L/min, fragmentor voltage 380 V, sheath gas temperature 350°C and flow 12 L/min, nozzle voltage 0 V, and capillary voltage -2000 V. Acquisition was set over the full *m/z* range 50-1000 at the MS acquisition rate of 2 spectra/s. All ion fragmentation (AIF) MS/MS analysis was performed on pooled QC samples (collision energy 0, 10, 30 eV).

An *in-house* database containing 600 polar metabolites was employed for metabolite annotation, using the Profinder B.08.00 software (Agilent Technologies) and targeted data mining in isotopologue extraction mode. The METLIN standard spectral library was used to validate putative metabolite identity via the MS/MS fragmentation pattern^51^ and EICs were used for relative metabolite quantification. Natural isotype abundance was corrected in ProFinder^52^. All tables and peak areas of detected metabolites were exported to R (http://cran.r-project.org/) and signal intensity drift correction was performed with the LOWESS/Spline normalization algorithm^53^ followed by noise filtering (CV pooled sample > 30%).

^13^C enrichment was calculated by taking into account the relative isotopologue abundances in two or more conditions as previously described^54,55^. Statistical significance was evaluated by applying univariate analysis on log10-transformed data, using a p value set arbitrarily to 0.05, followed by false discovery rate correction (Benjamini-Hochberg).

### Statistical analysis

Statistics are presented as mean ± SD from one representative dataset from at least three biological replicates, unless otherwise indicated. Statistical significance was calculated by Student’s t-test or ANOVA as specified in the figure legends, and exact p values are shown when lower than p<0.05. All graphs were generated using Prism 10 (GraphPad Software).

## References

1. Smith, R.L., Soeters, M.R., Wust, R.C.I., and Houtkooper, R.H. (2018). Metabolic Flexibility as an Adaptation to Energy Resources and Requirements in Health and Disease. Endocr Rev 39, 489–517. 10.1210/er.2017-00211.

2. Wishart, D.S., Guo, A., Oler, E., Wang, F., Anjum, A., Peters, H., Dizon, R., Sayeeda, Z., Tian, S., Lee, B.L., et al. (2022). HMDB 5.0: the Human Metabolome Database for 2022. Nucleic Acids Res 50, D622–D631. 10.1093/nar/gkab1062.

3. McMenamy, R.H., Lund, C.C., and Oncley, J.L. (1957). Unbound amino acid concentrations in human blood plasmas. J Clin Invest 36, 1672–1679. 10.1172/JCI103568.

4. Eagle, H., Oyama, V.I., Levy, M., Horton, C.L., and Fleischman, R. (1956). The growth response of mammalian cells in tissue culture to L-glutamine and L-glutamic acid. J Biol Chem 218, 607–616.

5. Zhang, J., Pavlova, N.N., and Thompson, C.B. (2017). Cancer cell metabolism: the essential role of the nonessential amino acid, glutamine. EMBO J 36, 1302–1315. 10.15252/embj.201696151.

6. Bott, A.J., Peng, I.C., Fan, Y., Faubert, B., Zhao, L., Li, J., Neidler, S., Sun, Y., Jaber, N., Krokowski, D., et al. (2015). Oncogenic Myc Induces Expression of Glutamine Synthetase through Promoter Demethylation. Cell Metab 22, 1068–1077. 10.1016/j.cmet.2015.09.025.

7. Wise, D.R., DeBerardinis, R.J., Mancuso, A., Sayed, N., Zhang, X.Y., Pfeiffer, H.K., Nissim, I., Daikhin, E., Yudkoff, M., McMahon, S.B., and Thompson, C.B. (2008). Myc regulates a transcriptional program that stimulates mitochondrial glutaminolysis and leads to glutamine addiction. Proc Natl Acad Sci U S A 105, 18782–18787. 10.1073/pnas.0810199105.

8. Zhao, X., Petrashen, A.P., Sanders, J.A., Peterson, A.L., and Sedivy, J.M. (2019). SLC1A5 glutamine transporter is a target of MYC and mediates reduced mTORC1 signaling and increased fatty acid oxidation in long-lived Myc hypomorphic mice. Aging Cell 18, e12947. 10.1111/acel.12947.

9. Yang, W.H., Qiu, Y., Stamatatos, O., Janowitz, T., and Lukey, M.J. (2021). Enhancing the Efficacy of Glutamine Metabolism Inhibitors in Cancer Therapy. Trends Cancer 7, 790–804. 10.1016/j.trecan.2021.04.003.

10. Cheng, T., Sudderth, J., Yang, C., Mullen, A.R., Jin, E.S., Mates, J.M., and DeBerardinis, R.J. (2011). Pyruvate carboxylase is required for glutamine-independent growth of tumor cells. Proc Natl Acad Sci U S A 108, 8674–8679. 10.1073/pnas.1016627108.

11. Pavlova, N.N., Hui, S., Ghergurovich, J.M., Fan, J., Intlekofer, A.M., White, R.M., Rabinowitz, J.D., Thompson, C.B., and Zhang, J. (2018). As Extracellular Glutamine Levels Decline, Asparagine Becomes an Essential Amino Acid. Cell Metab 27, 428–438 e425. 10.1016/j.cmet.2017.12.006.

12. Zhang, J., Fan, J., Venneti, S., Cross, J.R., Takagi, T., Bhinder, B., Djaballah, H., Kanai, M., Cheng, E.H., Judkins, A.R., et al. (2014). Asparagine plays a critical role in regulating cellular adaptation to glutamine depletion. Mol Cell 56, 205–218. 10.1016/j.molcel.2014.08.018.

13. Linder, S.J., Bernasocchi, T., Martinez-Pastor, B., Sullivan, K.D., Galbraith, M.D., Lewis, C.A., Ferrer, C.M., Boon, R., Silveira, G.G., Cho, H.M., et al. (2023). Inhibition of the proline metabolism rate-limiting enzyme P5CS allows proliferation of glutamine-restricted cancer cells. Nat Metab 5, 2131–2147. 10.1038/s42255-023-00919-3.

14. Alkan, H.F., Walter, K.E., Luengo, A., Madreiter-Sokolowski, C.T., Stryeck, S., Lau, A.N., Al-Zoughbi, W., Lewis, C.A., Thomas, C.J., Hoefler, G., et al. (2018). Cytosolic Aspartate Availability Determines Cell Survival When Glutamine Is Limiting. Cell Metab 28, 706–720 e706. 10.1016/j.cmet.2018.07.021.

15. Altea-Manzano, P., Cuadros, A.M., Broadfield, L.A., and Fendt, S.M. (2020). Nutrient metabolism and cancer in the in vivo context: a metabolic game of give and take. EMBO Rep 21, e50635. 10.15252/embr.202050635.

16. Birsoy, K., Wang, T., Chen, W.W., Freinkman, E., Abu-Remaileh, M., and Sabatini, D.M. (2015). An Essential Role of the Mitochondrial Electron Transport Chain in Cell Proliferation Is to Enable Aspartate Synthesis. Cell 162, 540–551. 10.1016/j.cell.2015.07.016.

17. Titov, D.V., Cracan, V., Goodman, R.P., Peng, J., Grabarek, Z., and Mootha, V.K. (2016). Complementation of mitochondrial electron transport chain by manipulation of the NAD+/NADH ratio. Science 352, 231–235. 10.1126/science.aad4017.

18. To, T.L., Cuadros, A.M., Shah, H., Hung, W.H.W., Li, Y., Kim, S.H., Rubin, D.H.F., Boe, R.H., Rath, S., Eaton, J.K., et al. (2019). A Compendium of Genetic Modifiers of Mitochondrial Dysfunction Reveals Intra-organelle Buffering. Cell 179, 1222–1238 e1217. 10.1016/j.cell.2019.10.032.

19. Yada, M., Hatakeyama, S., Kamura, T., Nishiyama, M., Tsunematsu, R., Imaki, H., Ishida, N., Okumura, F., Nakayama, K., and Nakayama, K.I. (2004). Phosphorylation-dependent degradation of c-Myc is mediated by the F-box protein Fbw7. EMBO J 23, 2116–2125. 10.1038/sj.emboj.7600217.

20. Koepp, D.M., Schaefer, L.K., Ye, X., Keyomarsi, K., Chu, C., Harper, J.W., and Elledge, S.J. (2001). Phosphorylation-dependent ubiquitination of cyclin E by the SCFFbw7 ubiquitin ligase. Science 294, 173–177. 10.1126/science.1065203.

21. Qi, Y., Rezaeian, A.H., Wang, J., Huang, D., Chen, H., Inuzuka, H., and Wei, W. (2024). Molecular insights and clinical implications for the tumor suppressor role of SCF(FBXW7) E3 ubiquitin ligase. Biochim Biophys Acta Rev Cancer 1879, 189140. 10.1016/j.bbcan.2024.189140.

22. Wei, W., Jin, J., Schlisio, S., Harper, J.W., and Kaelin, W.G., Jr. (2005). The v-Jun point mutation allows c-Jun to escape GSK3-dependent recognition and destruction by the Fbw7 ubiquitin ligase. Cancer Cell 8, 25–33. 10.1016/j.ccr.2005.06.005.

23. Welcker, M., Orian, A., Grim, J.E., Eisenman, R.N., and Clurman, B.E. (2004). A nucleolar isoform of the Fbw7 ubiquitin ligase regulates c-Myc and cell size. Curr Biol 14, 1852–1857. 10.1016/j.cub.2004.09.083.

24. Matsumoto, A., Tateishi, Y., Onoyama, I., Okita, Y., Nakayama, K., and Nakayama, K.I. (2011). Fbxw7beta resides in the endoplasmic reticulum membrane and protects cells from oxidative stress. Cancer Sci 102, 749–755. 10.1111/j.1349-7006.2011.01851.x.

25. Utter, M.F., and Keech, D.B. (1960). Formation of oxaloacetate from pyruvate and carbon dioxide. J Biol Chem 235, PC17–18.

26. Liano-Pons, J., Arsenian-Henriksson, M., and Leon, J. (2021). The Multiple Faces of MNT and Its Role as a MYC Modulator. Cancers (Basel) 13. 10.3390/cancers13184682.

27. Tsherniak, A., Vazquez, F., Montgomery, P.G., Weir, B.A., Kryukov, G., Cowley, G.S., Gill, S., Harrington, W.F., Pantel, S., Krill-Burger, J.M., et al. (2017). Defining a Cancer Dependency Map. Cell 170, 564–576 e516. 10.1016/j.cell.2017.06.010.

28. Saunders, A., Huang, X., Fidalgo, M., Reimer, M.H., Jr., Faiola, F., Ding, J., Sanchez-Priego, C., Guallar, D., Saenz, C., Li, D., and Wang, J. (2017). The SIN3A/HDAC Corepressor Complex Functionally Cooperates with NANOG to Promote Pluripotency. Cell Rep 18, 1713–1726. 10.1016/j.celrep.2017.01.055.

29. Lavin, D.P., Abassi, L., Inayatullah, M., and Tiwari, V.K. (2021). Mnt Represses Epithelial Identity To Promote Epithelial-to-Mesenchymal Transition. Mol Cell Biol 41, e0018321. 10.1128/MCB.00183-21.

30. Thirimanne, H.N., Wu, F., Janssens, D.H., Swanger, J., Diab, A., Feldman, H.M., Amezquita, R.A., Gottardo, R., Paddison, P.J., Henikoff, S., and Clurman, B.E. (2022). Global and context-specific transcriptional consequences of oncogenic Fbw7 mutations. Elife 11. 10.7554/eLife.74338.

31. Zhang, J., Lee, D., Dhiman, V., Jiang, P., Xu, J., McGillivray, P., Yang, H., Liu, J., Meyerson, W., Clarke, D., et al. (2020). An integrative ENCODE resource for cancer genomics. Nat Commun 11, 3696. 10.1038/s41467-020-14743-w.

32. Sullivan, M.R., Danai, L.V., Lewis, C.A., Chan, S.H., Gui, D.Y., Kunchok, T., Dennstedt, E.A., Vander Heiden, M.G., and Muir, A. (2019). Quantification of microenvironmental metabolites in murine cancers reveals determinants of tumor nutrient availability. Elife 8. 10.7554/eLife.44235.

33. Akhoondi, S., Sun, D., von der Lehr, N., Apostolidou, S., Klotz, K., Maljukova, A., Cepeda, D., Fiegl, H., Dafou, D., Marth, C., et al. (2007). FBXW7/hCDC4 is a general tumor suppressor in human cancer. Cancer Res 67, 9006–9012. 10.1158/0008-5472.CAN-07-1320.

34. Tate, J.G., Bamford, S., Jubb, H.C., Sondka, Z., Beare, D.M., Bindal, N., Boutselakis, H., Cole, C.G., Creatore, C., Dawson, E., et al. (2019). COSMIC: the Catalogue Of Somatic Mutations In Cancer. Nucleic Acids Res 47, D941–D947. 10.1093/nar/gky1015.

35. Davis, R.J., Gonen, M., Margineantu, D.H., Handeli, S., Swanger, J., Hoellerbauer, P., Paddison, P.J., Gu, H., Raftery, D., Grim, J.E., et al. (2018). Pan-cancer transcriptional signatures predictive of oncogenic mutations reveal that Fbw7 regulates cancer cell oxidative metabolism. Proc Natl Acad Sci U S A 115, 5462–5467. 10.1073/pnas.1718338115.

36. Xia, H., Dufour, C.R., Medkour, Y., Scholtes, C., Chen, Y., Guluzian, C., B’Chir, W., and Giguere, V. (2023). Hepatocyte FBXW7-dependent activity of nutrient-sensing nuclear receptors controls systemic energy homeostasis and NASH progression in male mice. Nat Commun 14, 6982. 10.1038/s41467-023-42785-3.

37. Sullivan, L.B., Gui, D.Y., Hosios, A.M., Bush, L.N., Freinkman, E., and Vander Heiden, M.G. (2015). Supporting Aspartate Biosynthesis Is an Essential Function of Respiration in Proliferating Cells. Cell 162, 552–563. 10.1016/j.cell.2015.07.017.

38. Sanjana, N.E., Shalem, O., and Zhang, F. (2014). Improved vectors and genome-wide libraries for CRISPR screening. Nat Methods 11, 783–784. 10.1038/nmeth.3047.

39. Sanson, K.R., Hanna, R.E., Hegde, M., Donovan, K.F., Strand, C., Sullender, M.E., Vaimberg, E.W., Goodale, A., Root, D.E., Piccioni, F., and Doench, J.G. (2018). Optimized libraries for CRISPR-Cas9 genetic screens with multiple modalities. Nat Commun 9, 5416. 10.1038/s41467-018-07901-8.

40. Skinner, O.S., Blanco-Fernandez, J., Goodman, R.P., Kawakami, A., Shen, H., Kemeny, L.V., Joesch-Cohen, L., Rees, M.G., Roth, J.A., Fisher, D.E., et al. (2023). Salvage of ribose from uridine or RNA supports glycolysis in nutrient-limited conditions. Nat Metab 5, 765–776. 10.1038/s42255-023-00774-2.

41. Kulak, N.A., Pichler, G., Paron, I., Nagaraj, N., and Mann, M. (2014). Minimal, encapsulated proteomic-sample processing applied to copy-number estimation in eukaryotic cells. Nat Methods 11, 319–324. 10.1038/nmeth.2834.

42. Wisniewski, J.R., and Gaugaz, F.Z. (2015). Fast and sensitive total protein and Peptide assays for proteomic analysis. Anal Chem 87, 4110–4116. 10.1021/ac504689z.

43. Meier, F., Brunner, A.D., Koch, S., Koch, H., Lubeck, M., Krause, M., Goedecke, N., Decker, J., Kosinski, T., Park, M.A., et al. (2018). Online Parallel Accumulation-Serial Fragmentation (PASEF) with a Novel Trapped Ion Mobility Mass Spectrometer. Mol Cell Proteomics 17, 2534–2545. 10.1074/mcp.TIR118.000900.

44. Meier, F., Brunner, A.D., Frank, M., Ha, A., Bludau, I., Voytik, E., Kaspar-Schoenefeld, S., Lubeck, M., Raether, O., Bache, N., et al. (2020). diaPASEF: parallel accumulation-serial fragmentation combined with data-independent acquisition. Nat Methods 17, 1229–1236. 10.1038/s41592-020-00998-0.

45. Cox, J., Hein, M.Y., Luber, C.A., Paron, I., Nagaraj, N., and Mann, M. (2014). Accurate proteome-wide label-free quantification by delayed normalization and maximal peptide ratio extraction, termed MaxLFQ. Mol Cell Proteomics 13, 2513–2526. 10.1074/mcp.M113.031591.

46. Tyanova, S., Temu, T., Sinitcyn, P., Carlson, A., Hein, M.Y., Geiger, T., Mann, M., and Cox, J. (2016). The Perseus computational platform for comprehensive analysis of (prote)omics data. Nat Methods 13, 731–740. 10.1038/nmeth.3901.

47. Medina, J., van der Velpen, V., Teav, T., Guitton, Y., Gallart-Ayala, H., and Ivanisevic, J. (2020). Single-Step Extraction Coupled with Targeted HILIC-MS/MS Approach for Comprehensive Analysis of Human Plasma Lipidome and Polar Metabolome. Metabolites 10. 10.3390/metabo10120495.

48. van der Velpen, V., Teav, T., Gallart-Ayala, H., Mehl, F., Konz, I., Clark, C., Oikonomidi, A., Peyratout, G., Henry, H., Delorenzi, M., et al. (2019). Systemic and central nervous system metabolic alterations in Alzheimer’s disease. Alzheimers Res Ther 11, 93. 10.1186/s13195-019-0551-7.

49. Gallart-Ayala, H., Konz, I., Mehl, F., Teav, T., Oikonomidi, A., Peyratout, G., van der Velpen, V., Popp, J., and Ivanisevic, J. (2018). A global HILIC-MS approach to measure polar human cerebrospinal fluid metabolome: Exploring gender-associated variation in a cohort of elderly cognitively healthy subjects. Anal Chim Acta 1037, 327–337. 10.1016/j.aca.2018.04.002.

50. Pang, Z., Lu, Y., Zhou, G., Hui, F., Xu, L., Viau, C., Spigelman, A.F., MacDonald, P.E., Wishart, D.S., Li, S., and Xia, J. (2024). MetaboAnalyst 6.0: towards a unified platform for metabolomics data processing, analysis and interpretation. Nucleic Acids Res 52, W398–W406. 10.1093/nar/gkae253.

51. Guijas, C., Montenegro-Burke, J.R., Domingo-Almenara, X., Palermo, A., Warth, B., Hermann, G., Koellensperger, G., Huan, T., Uritboonthai, W., Aisporna, A.E., et al. (2018). METLIN: A Technology Platform for Identifying Knowns and Unknowns. Anal Chem 90, 3156–3164. 10.1021/acs.analchem.7b04424.

52. Midani, F.S., Wynn, M.L., and Schnell, S. (2017). The importance of accurately correcting for the natural abundance of stable isotopes. Anal Biochem 520, 27–43. 10.1016/j.ab.2016.12.011.

53. Tsugawa, H., Kanazawa, M., Ogiwara, A., and Arita, M. (2014). MRMPROBS suite for metabolomics using large-scale MRM assays. Bioinformatics 30, 2379–2380. 10.1093/bioinformatics/btu203.

54. Roci, I., Gallart-Ayala, H., Schmidt, A., Watrous, J., Jain, M., Wheelock, C.E., and Nilsson, R. (2016). Metabolite Profiling and Stable Isotope Tracing in Sorted Subpopulations of Mammalian Cells. Anal Chem 88, 2707–2713. 10.1021/acs.analchem.5b04071.

55. Jain, M., Nilsson, R., Sharma, S., Madhusudhan, N., Kitami, T., Souza, A.L., Kafri, R., Kirschner, M.W., Clish, C.B., and Mootha, V.K. (2012). Metabolite profiling identifies a key role for glycine in rapid cancer cell proliferation. Science 336, 1040–1044. 10.1126/science.1218595.

